## Supplementary Files for "Widespread exposure to SARS-CoV-2 in wildlife communities"

Supplementary Information Text

**Site descriptions**

For all sites, traps were placed near structures (homes, sheds, and fences) as well as in and around vegetation (trees, bushes, gardens). We had two urban sites located in Blacksburg, VA (BB) and Roanoke, VA (RK). All homes were located in residential neighborhoods composed of single family or multiunit houses. Our locations in Roanoke were all on public land located in three small urban parks along the Roanoke River Greenway or on Mill Mountain Park. The small parks are located along a major bike/walking paved trail that runs along a river through the center of Roanoke. These parks were all located in mixed-use areas that included nearby houses, apartment buildings, hospitals, and businesses. Preston Forest (PF) is an intermediary site between urban and rural. Houses are spaced further apart (~5 acre lots) and hence the human population is at a lower density. Furthermore, there are no sidewalks so foot traffic is reduced. We trapped in the backyards of two homes that are mainly dominated by woody vegetation and leaf debris.

New River Trails State Park (Foster Falls; NRT) is a section of a larger State Park located along the New River. Foster Falls has a campground and is a popular area for outdoor recreation. Brush Mountain (BM) is located within a complex of private, non-profit owned lands that are open to the public for hiking and biking. The area we trapped was not yet open to the public as it was in the process of being prepared (building a parking area, grooming trails, etc). Mountain Lake Biological Station site (ML) is a research field station with a rotating group of people using the facilities, including classrooms and housing. We trapped away from the housing complex in the surrounding forest. There was a hiking trail located adjacent to our sites and a commercial lodge about two miles down the road. However, the lands owned by the research station, where we trapped, were off-limits to public use. Pandapas Pond (PP) is located in Montgomery county and is a high use area for outdoor recreation including trail running, hiking, mountain biking, and fishing. We placed traps along a forest service road. The area is dominated by trees and leaf debris on the ground. Caldwell fields (CF) is located also in Montgomery county. The area is lightly used for general outdoor recreation including hunting.

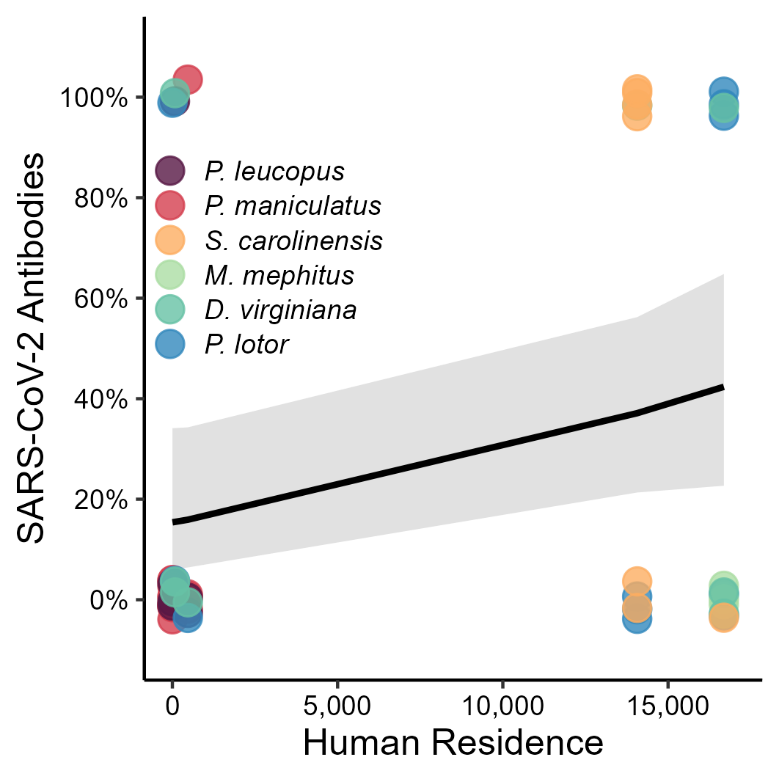

**Supplementary Figure 1.** **Examination of SARS-CoV-2 exposure relationship between human residence (U.S. 2020 census data) and seroprevalence collected from 5 different sites in VA, USA.** Grey ribbons (shadowed area) represent 95% confidence intervals (intercept = -1.277, β = 0.607, P = 0.059).

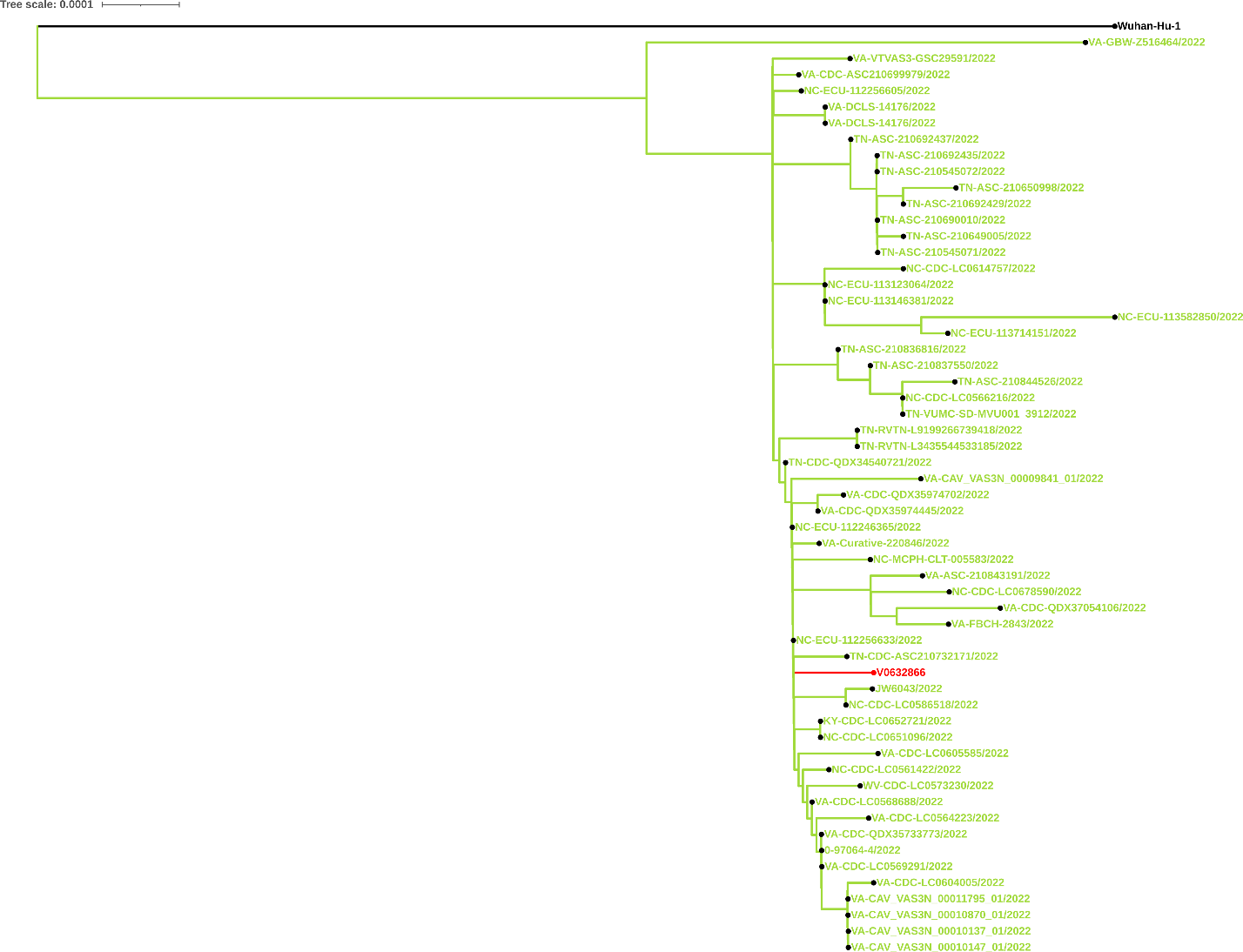
 **Supplementary Figure 2.** Phylogenetic tree for a Virginia opossum (V0652866) assigned to the BA.2.10.1 Pango lineage. See Supplementary Table 11 for further information. Red indicates the wildlife sample.

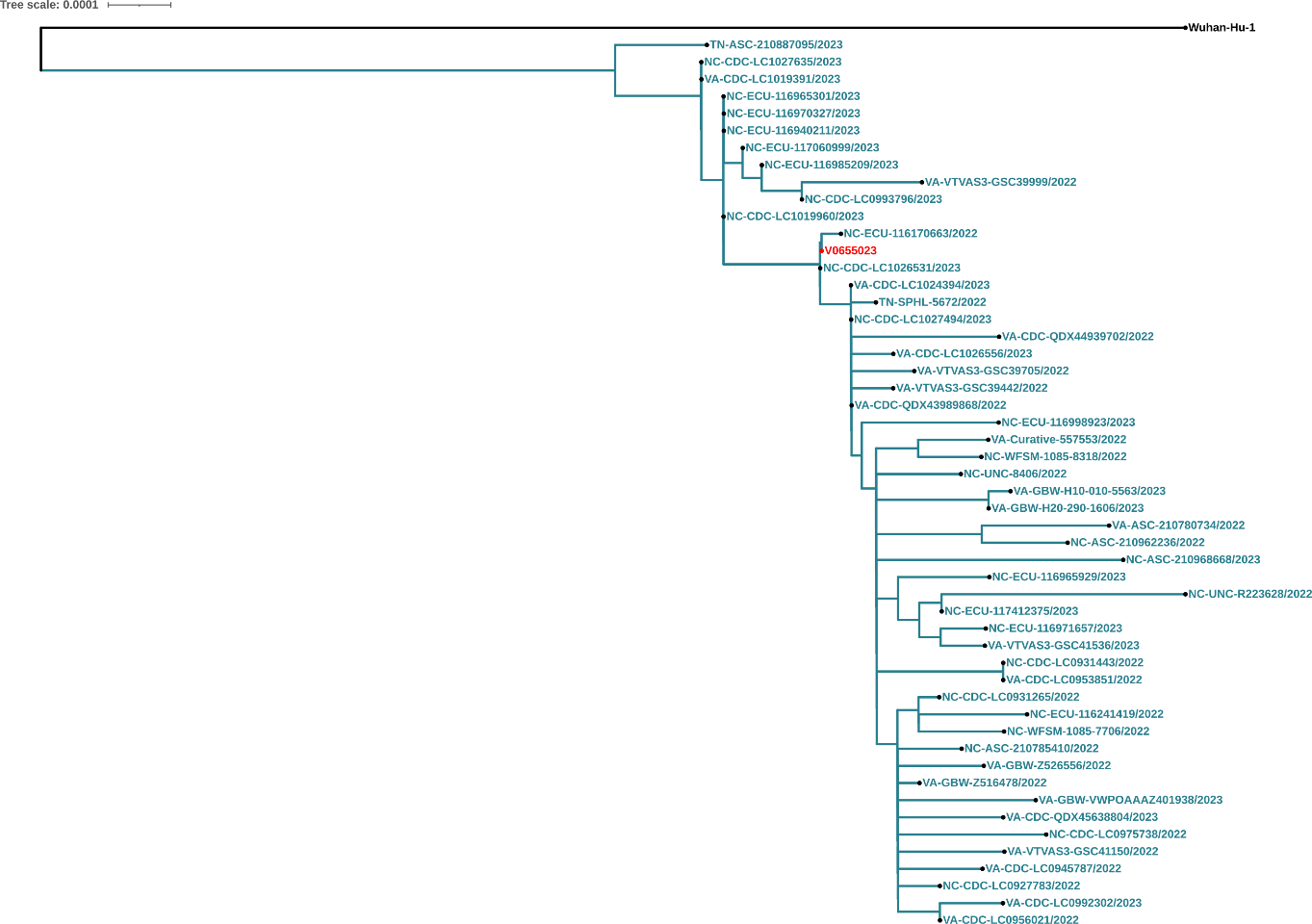

**Supplementary Figure 3.** Phylogenetic tree for a deer mouse (V0655023) assigned to the XBB Pango lineage. See Supplementary Table 11 for further information. Red indicates the wildlife sample.

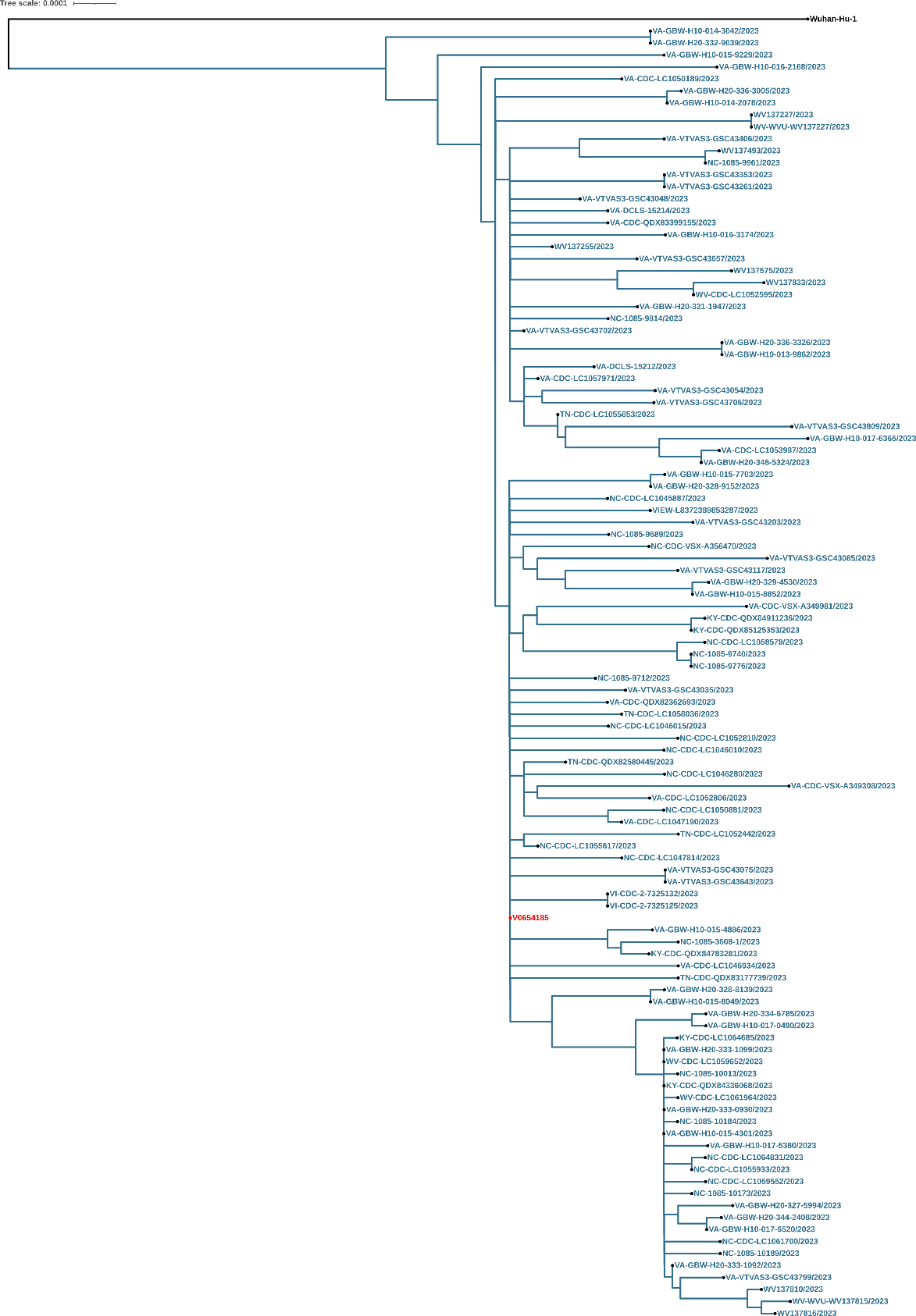

**Supplementary Figure 4.** Phylogenetic tree for a raccoon (V0654185) assigned to the XBB.1.5 Pango lineage. See Supplementary Table 11 for further information. Red indicates the wildlife sample.

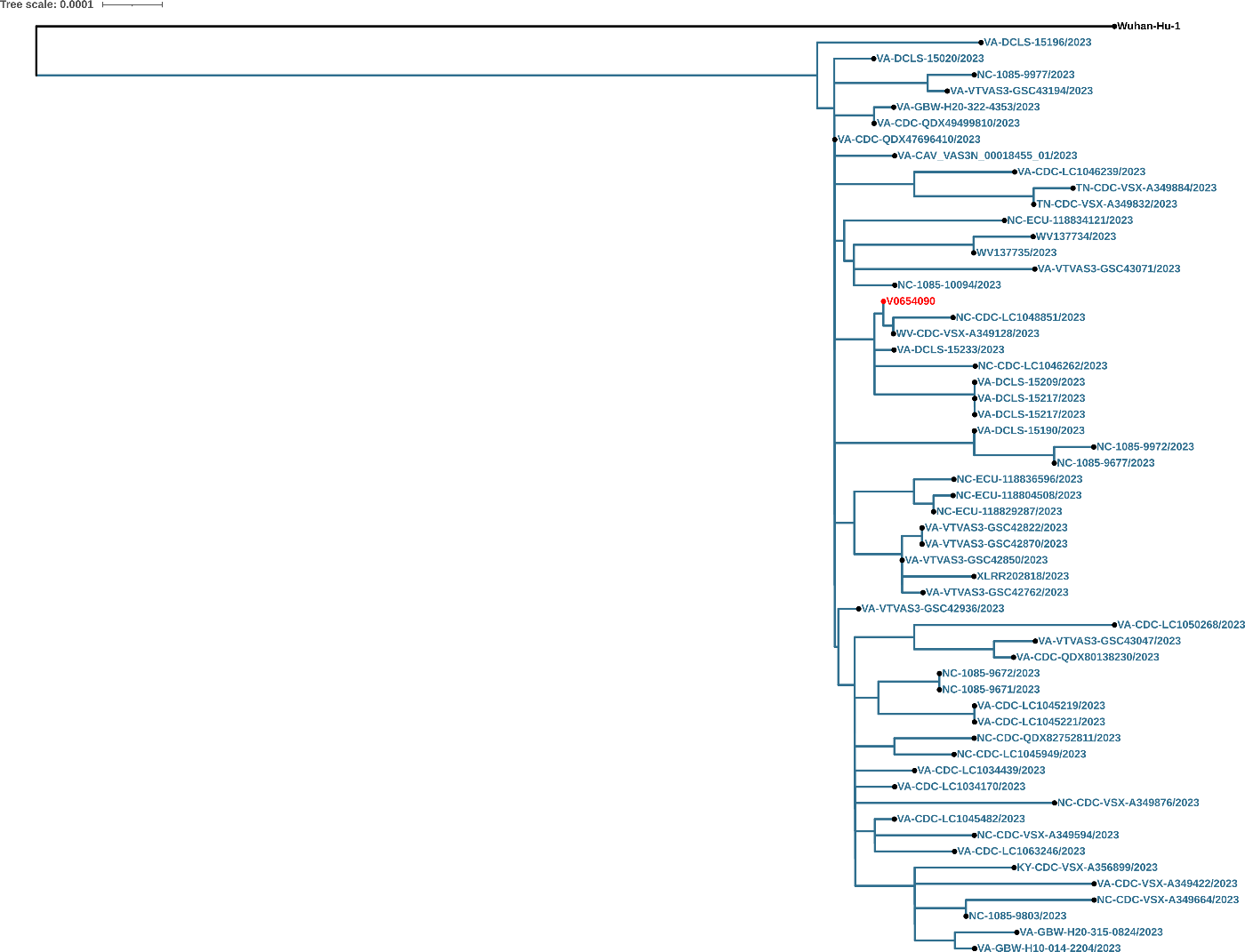

**Supplementary Figure 5.** Phylogenetic tree for a Virginia opossum (V0654090) assigned to the XBB.1.5.10 Pango lineage. See Supplementary Table 11 for further information. Red indicates the wildlife sample.

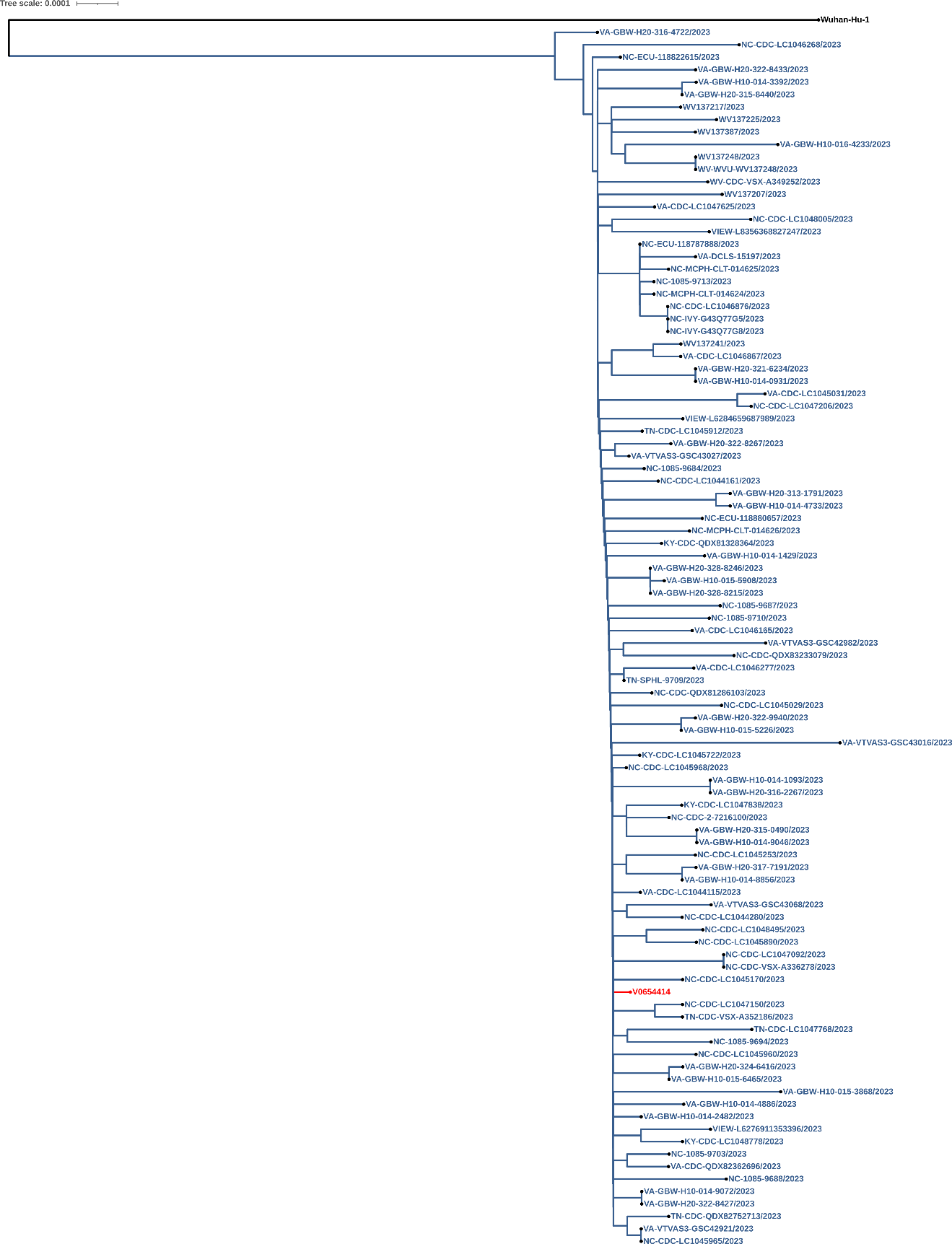

**Supplementary Figure 6.** Phylogenetic tree for an Eastern cottontail (V0654414) assigned to the XBB.1.16 Pango lineage. See Supplementary Table 11 for further information. Red indicates the wildlife sample.

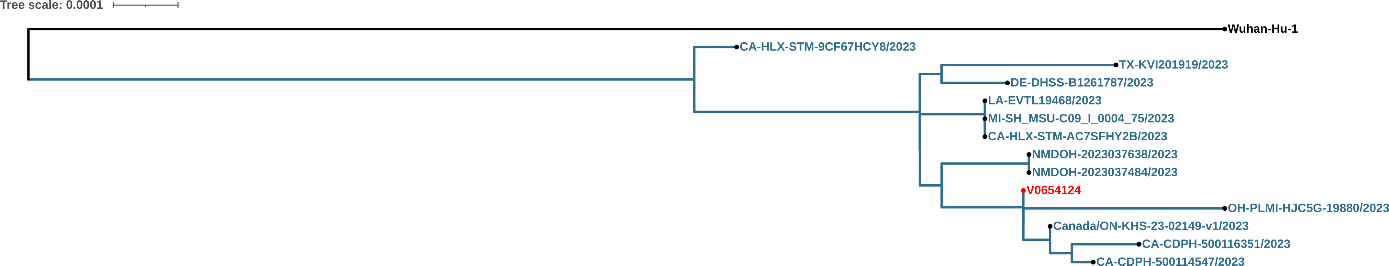

**Supplementary Figure 7.** Phylogenetic tree for a groundhog (V0654124) assigned to XBB.1.5.45 Pango lineage. See Supplementary Table 11 for further information. Red indicates the wildlife sample.

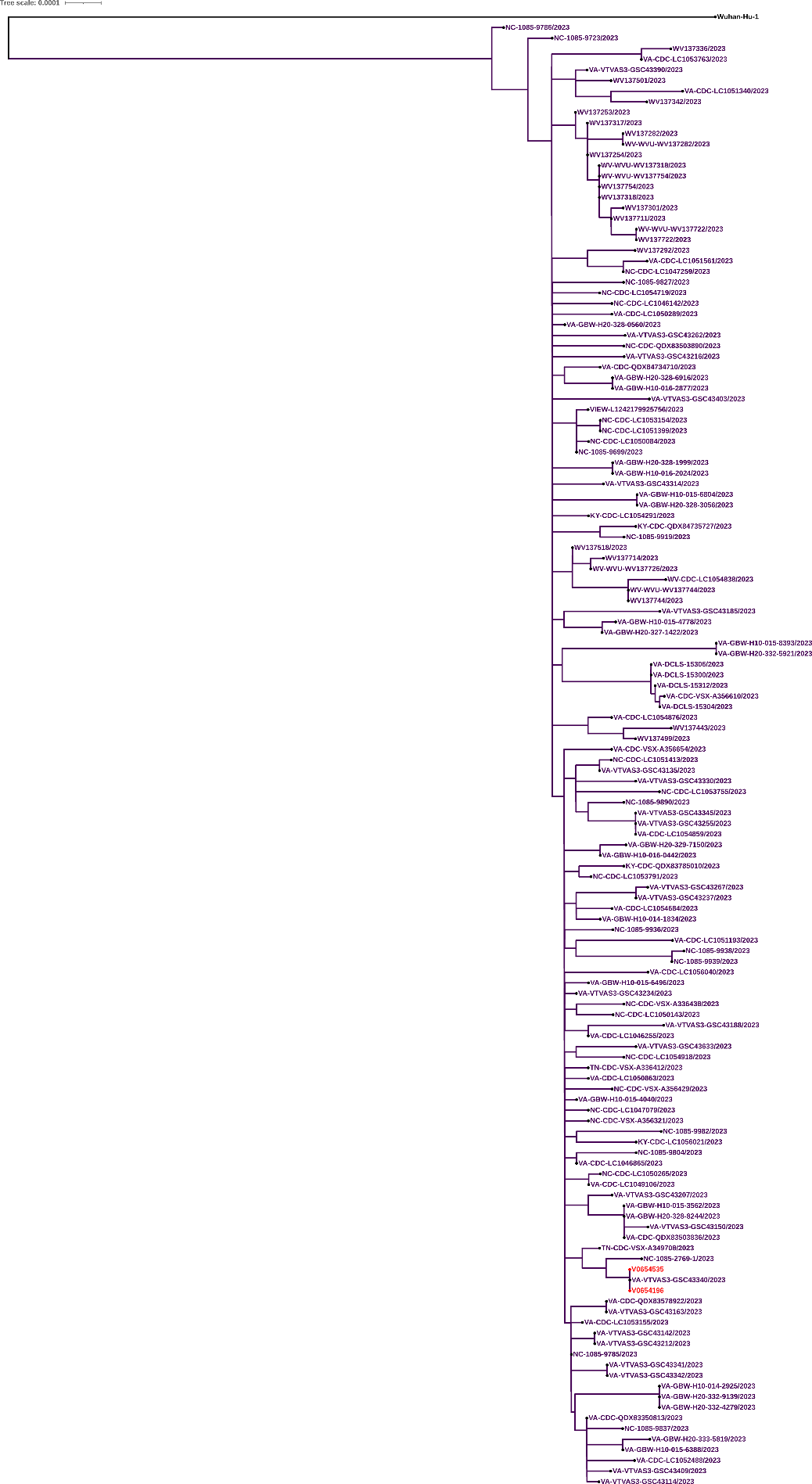
**Supplementary Figure 8.** Phylogenetic tree for two deer mice (V0654196 and V0653535) assigned to the EG.5.1.1 Pango lineage. See Supplementary Table 11 for further information. Red indicates the wildlife sample.

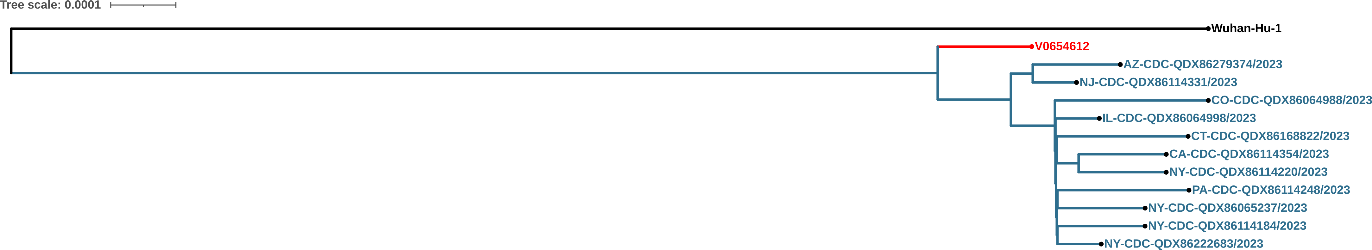

**Supplementary Figure 9.** Phylogenetic tree for a deer mouse (V0654612) assigned to the JD.1 Pango lineage. See Supplementary Table 11 for further information. Red indicates the wildlife sample.

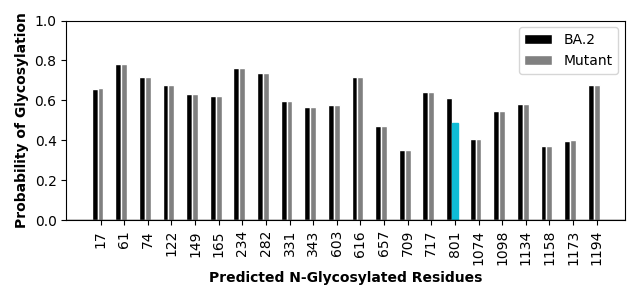
 **Supplementary Figure 10.** Predicted N-glycosylated residues identified by the NetNGlyc 1.0 server with the probability of being glycosylated based on BA.2 or [mutant] sequence.

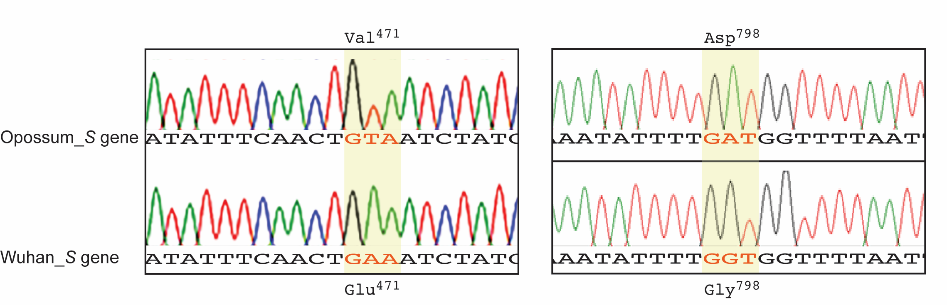

**Supplementary Figure 11. Sanger sequencing peaks of the genomic regions for the positive opossum from July 2022.** Flanking the Glu^471^Val (left) and Gly^798^Asp (right) mutations identified in the *S* gene of the opossum-infected SARS-CoV-2 sample and its corresponding sequence in the original Wuhan strain. Nucleotide mutations and matching wild-type sequences are shaded in yellow.

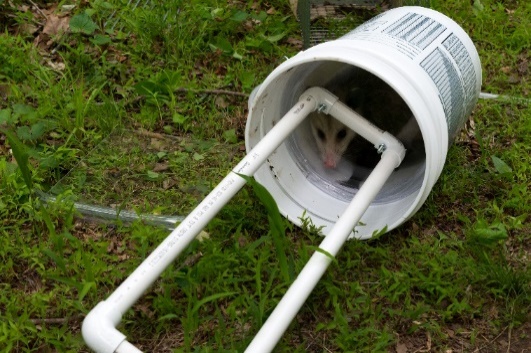

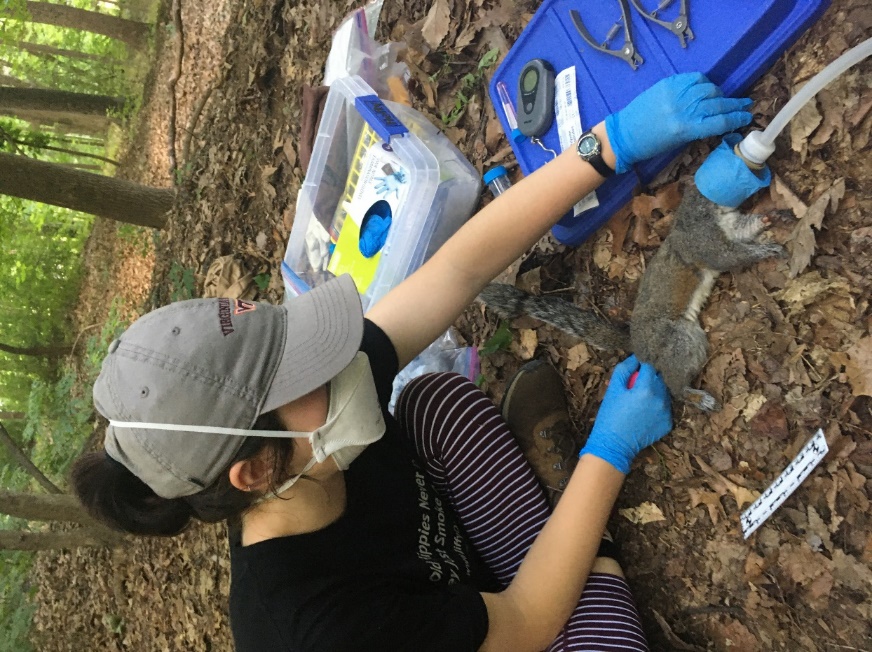

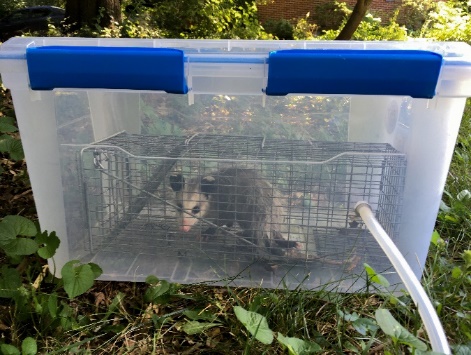

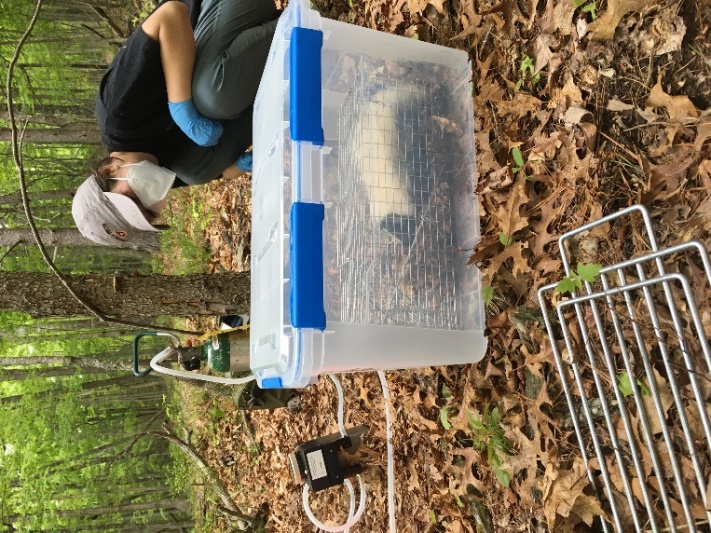

**a**

**b**

**c**

**d**

**Supplementary Figure 12. Equipment modified for animal processing.** A modified plastic container (a & b). We used a vaporizer with a small O2 tank to supply isoflurane into the chambers (b). We made masks from the top part of a plastic bottle (c). We used a modified bucket chamber for anesthetizing animals trapped in larger cages (d).

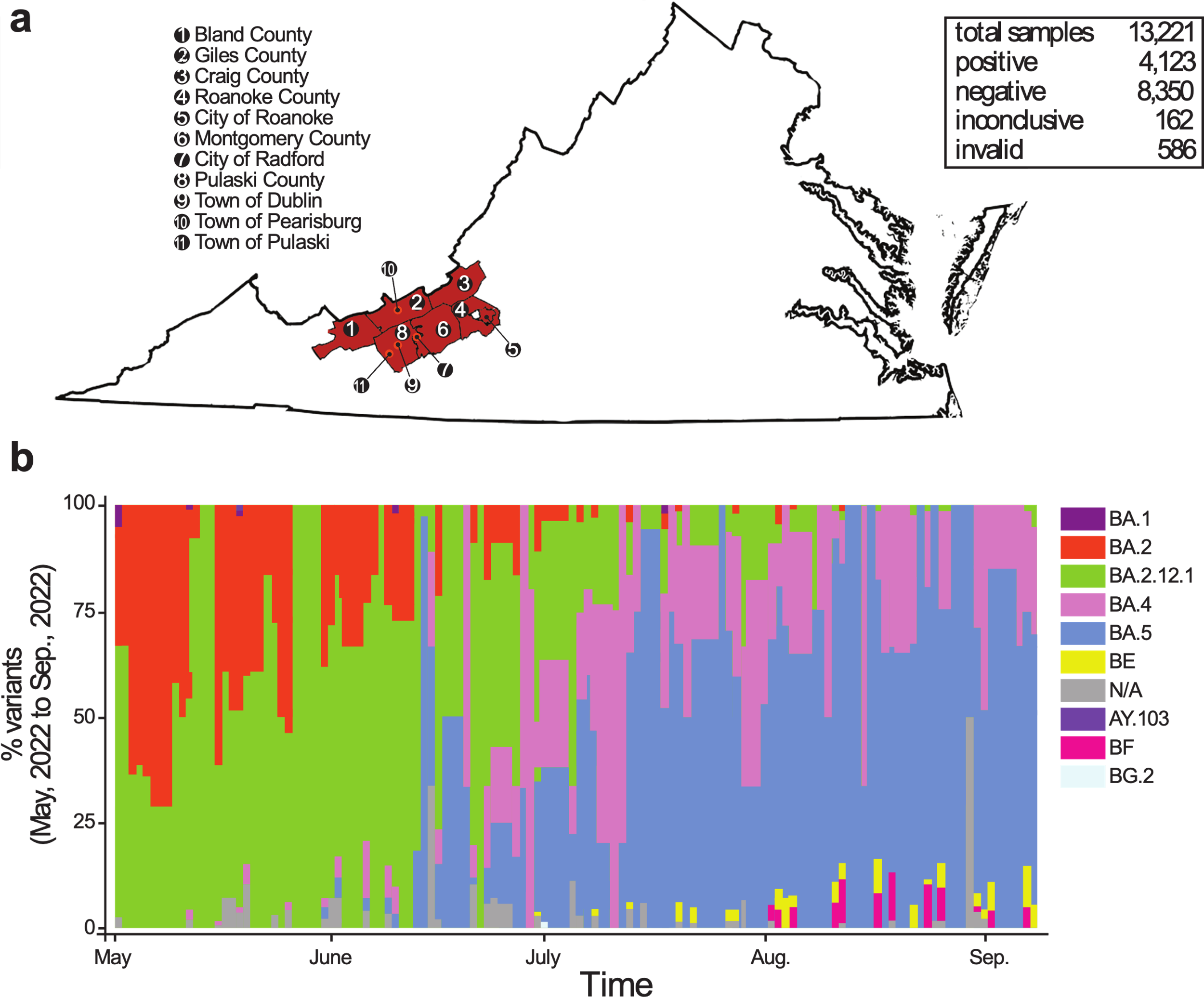

**Supplementary Figure 13. Distribution of SARS-CoV-2 variants in circulation during the 2022 sampling period.** (a) Map of counties where sequence data was obtained. (b) Summary of weekly distribution of SARS-CoV-2 variants circulating in human communities in Southwest Virginia between May and September, 2022 as determined by RT-qPCR, RMA, and WGS sequencing.

Supplementary Table 1. Summary of species tested including both RT-qPCR and serology (60% cut-off) results for SARS-CoV-2 after the virus had arrived in the United States.

| Species | RT-qPCR (n) | RT-qPCR positive ≥2 genes^1^ | RT-qPCR positive 1 gene^1^ | Number of WGS obtained | Number of partial WGS obtained | Serology (n) | Sero-positive samples |
| --- | --- | --- | --- | --- | --- | --- | --- |
| *Peromyscus maniculatus^2^* | 172 | 8 (4N+E+S, 1N+E, 1N+S, 2E+S) | 9 (4N, 1E, 4S) | 4 | 1 | 14 | 1 |
| *Procyon lotor^2^* | 84 | 4 (3N+E+S, 1N+S) | 5 (2N, 3S) | 1 |  | 11 | 4 |
| *Didelphis virginiana^2^* | 140 | 4 (1N+S+E, 1N+S, 1N+E, 1E+S) | 16 (2N, 6E, 8S) | 2 |  | 8 | 2 |
| *Sciurus carolinensis^2^* | 105 |  | 4 (1E, 3S) |  |  | 7 | 4 |
| *Peromyscus leucopus^2^* | 19 |  |  |  |  | 6 | 0 |
| *Mephitis mephitis^2^* | 25 |  | 1 (1N) |  |  | 3 | 2 |
| *Vulpes vulpes^2^* | 17 |  | 2 (1N, 1S) |  |  |  |  |
| *Odocoileus virginianus* | 20 |  | 3 (3S) |  |  |  |  |
| *Lynx rufus* | 3 |  | 2 (2S) |  |  |  |  |
| *Sylvilagus floridanus* | 118 | 3 (1N+E+S, 1N+S, 1E+S) | 9 (7N, 1E 1S) | 1 |  |  |  |
| *Marmota monax^2^* | 31 | 3 (2N+E, 1E+S) | 2 (2N) | 1 |  |  |  |
| *Tamias striatus^2^* | 12 |  |  |  |  |  |  |
| *Lasiurus borealis^2^* | 12 | 1 (1N+S) | 1 (1N) | 0 | 2 |  |  |
| *Eptesicus fuscus^2^* | 7 |  |  |  |  |  |  |
| *Ursus americanus* | 7 |  | 1 (1N) |  |  |  |  |
| *Sciurus niger* | 3 |  |  |  |  |  |  |
| *Blarina brevicada^2^* | 3 |  |  |  |  |  |  |
| *Rattus rattus* | 2 |  |  |  |  |  |  |
| *Urocyon cinereoargenteus* | 2 |  |  |  |  |  |  |
| *Castor canadensis* | 2 |  | 1 (1S) |  |  |  |  |
| *Peromyscus sp.* | 2 |  |  |  |  |  |  |
| *Mustela vison* | 1 |  |  |  |  |  |  |
| *Microtus pennsylvanicus* | 1 |  |  |  |  |  |  |
| *Mus musculus* | 1 |  |  |  |  |  |  |
| Total | 789 | 23 | 53 | 9 | 3 | 49 | 24 |
| ^1^Letter denotes SARS-CoV-2 genes(s) that amplified.  *^2^*Species sampled at one of the 8 study sites | | | | | | | |

Supplementary Table 2. Average RT-qPCR values for all samples in Virginia and Washington DC from June 2022 through September 2023. Samples were collected either in the field (Study) or by a Virginia wildlife rehabilitation center (Blue Ridge Wildlife Center (BRWC), The Wildlife Center of Virginia (TWCV), or Southwest Virginia Wildlife Center of Roanoke (SWVAWC)). See attached excel file.

**Supplementary Table 3.** Comparative analysis of SARS-CoV-2 sequences from wild animals isolated in southwest Virginia. Consensus sequences for SARS-CoV-2 isolated from wild animal specimen were analyzed for sequence coverage and percent identity compared to the SARS-CoV-2 reference genome (NC_045512), Pango lineage, closely related SARS-CoV-2 sequence isolated from a human, and unique amino acid substitutions, which are not found in their closely related sequences. See Table S5 for unresolved sequences.

| Sample ID  (Accession number) | Animal species | Location | Date of isolation | Sequence coverage (Percent Identity) | Pango | Closely related sequence, state & collection date | Unique amino acid substitutions |
| --- | --- | --- | --- | --- | --- | --- | --- |
| V0632866  (OR866905) | *Didelphis* *virginiana* | Blacksburg, Montgomery Co, VA | 05/29/2022 | 96.8%  (99.8%) | BA.2.10.1 | OM999909  TN 03/06/2022 | ORF1a:T2495I  S:E471V |
| V0654090  (OR878666) | *Didelphis* *virginiana* | Star Tannery, Frederick Co, VA | 07/10/2023 | 31.8%  (99.0%) | XBB.1.5.10 | OR454205  VA 7/18/2023 | none |
| V0654124  (OR866349) | *Marmota monax* | Front Royal, Warren Co, VA | 06/30/2023 | 73.5%  (99.8%) | XBB.1.5.45 | EPI_ISL_17744393  OH 06/02/2023* | S:H146Q |
| V0654196  (OR866382) | *Peromyscus maniculatus* | Blacksburg, Montgomery Co, VA | 09/02/2023 | 99.1%  (99.5%) | EG.5.1.1 | EPI_ISL_18287981  VA 08/29/2023 | none |
| V0654535  (OR866443) | *Peromyscus maniculatus* | Blacksburg, Montgomery Co, VA | 09/02/2023 | 88.6%  (99.4%) | EG.5.1.1 | EPI_ISL_18287981  VA 08/29/2023 | none |
| V0654612  (OR878668) | *Peromyscus maniculatus* | Blacksburg, Montgomery Co, VA | 09/02/2023 | 24.3%  (99.1%) | JD.1 | OR708231  NJ 10/06/23 | none |
| V0655023  (OR866910) | *Peromyscus maniculatus* | Blacksburg, Montgomery Co, VA | 09/06/2023 | 59.7%  (99.8%) | XBB | EPI_ISL_16384582  NC 11/26/22 | none |
| V0654185  (OR878667) | *Procyon lotor* | Max Meadows, Wythe Co, VA | 09/06/2023 | 25.0%  (99.6%) | XBB.1.5 | EPI_ISL_18124823  VA 08/13/2023 | none |
| V0654414  (OR866437) | *Sylvilagus floridanus* | Stafford, Stafford Co, VA | 07/17/2023 | 82.1%  (99.8%) | XBB.1.16 | OR252035  NC 06/22/23 | none |

**Supplementary Table 4.** Summary of SARS-CoV-2 amplicon sequences obtained from wild animals. Sequences were analyzed for their length in bases and their position and percent identity relative to the reference genome (NC_045512.2).

| Sample ID (Accession number) | Animal Species | Location | Date | Sequence length | SARS-CoV-2 sequence position | Percent identity (mutation count) |
| --- | --- | --- | --- | --- | --- | --- |
| V0654648  (OR871756  OR872533) | *Lasiurus borealis* | Blacksburg, Montgomery Co, VA | 8/21/2023 | 272  267 | 24508..24779  28516..28782 | 100% (0)  100% (0) |
| V0655027  (OR871072) | *Lasiurus borealis* | Blacksburg, Montgomery Co, VA | 8/21/2023 | 249 | 24812..25060 | 99% (2) |
| V0654636  (OR871750  OR872518  OR871751) | *Peromyscus maniculatus* | Blacksburg, Montgomery Co, VA | 09/02/2023 | 377  247  317 | 23570..23946  28196..28451  28464..28780 | 99% (3)  95% (12)  100% (0) |

Supplementary Table 5. Serology results from the 49 samples collected after SARS-CoV-2 arrival (post) and 67 samples prior to SARS-CoV-2 arrival (pre) samples collected. All ‘pre’ *Peromyscus* species were provided by NEON^1^, the Eastwood lab at Virginia Tech, or the Kilpatrick lab at UC Santa Cruz^2^. All counties are located in Virginia unless otherwise noted. See attached excel file.

| **Supplementary Table 6.** Number of seropositive samples under 4 different percent neutralization cutoff values from the 49 samples collected in summer 2022 in Virginia, U.S.A from 6 species. The four different percent neutralization cutoff values we evaluated include: 40% (Pos 40), 50% (Pos 50), 60% (Pos 60; the value we used for Fig 2c), and 70% (Pos 70). Additionally, we include the seropositivity values for each species under the 4 different cutoff values. | | | | | | | | | |
| --- | --- | --- | --- | --- | --- | --- | --- | --- | --- |
| Species | Pos 40 | Pos 50 | Pos 60 | Pos 70 | N | Percent Pos 40 | Percent Pos 50 | Percent Pos 60 | Percent Pos 70 |
| *Peromyscus leucopus* | 2 | 1 | 1 | 1 | 6 | 33.33% | 16.67% | 16.67% | 16.67% |
| *Peromyscus maniculatus* | 6 | 4 | 1 | 0 | 14 | 42.86% | 28.57% | 7.14% | 0.00% |
| *Sciurus carolinensis* | 6 | 5 | 4 | 2 | 7 | 85.71% | 71.43% | 57.14% | 28.57% |
| *Mephitus mephitus* | 2 | 2 | 0 | 0 | 3 | 66.67% | 66.67% | 0.00% | 0.00% |
| *Didelphis* *virginiana* | 6 | 5 | 3 | 1 | 8 | 75.00% | 62.50% | 37.50% | 12.50% |
| *Procyon lotor* | 9 | 7 | 4 | 3 | 11 | 81.82% | 63.64% | 36.36% | 27.27% |
| Total | 31 | 24 | 13 | 7 | 49 | 63.27% | 48.98% | 26.53% | 14.29% |

| **Supplementary Table 7.** Generalized linear mixed model results evaluating the relationship between urbanization and seroprevalence of mammals with species as a random effect. We evaluated this relationship using 5 different % neutralization cutoff values. See Fig. 2 for predicted relationship from the 60% cutoff model. | | | | | | | |
| --- | --- | --- | --- | --- | --- | --- | --- |
| % Neutralization cutoff | Intercept | | |  | Imperviousness | | |
|  | β | SE | p |  | β | SE | p |
| 40% | -0.054 | 0.372 | 0.884 |  | 0.053 | 0.024 | 0.026 |
| 50% | -0.608 | 0.380 | 0.109 |  | 0.042 | 0.019 | 0.024 |
| 60% | -1.655 | 0.477 | 0.001 |  | 0.038 | 0.018 | 0.032 |
| 65% | -1.892 | 0.514 | 0.000 |  | 0.044 | 0.019 | 0.017 |
| 70% | -1.885 | 0.561 | 0.001 |  | 0.005 | 0.179 | 0.858 |

| **Supplementary Table 8.** Summary of serology and RT-qPCR results from samples collected in Virginia and Washington D.C. in 2022 and 2023. | | | | | | |
| --- | --- | --- | --- | --- | --- | --- |
| Species | Serology (n) | Sero-positive (Y/N) | RT-qPCR (n) | RT-qPCR positive (Y/N) | WGS (Y/N) | Partial Sequence (Y/N) |
| *Peromyscus maniculatus* | 14 | Y | 172 | Y | Y | Y |
| *Procyon lotor* | 11 | Y | 84 | Y | Y | N |
| *Didelphis virginiana* | 8 | Y | 140 | Y | Y | N |
| *Sciurus carolinensis* | 7 | Y | 105 | N | N | N |
| *Peromyscus leucopus* | 6 | N | 19 | N | N | N |
| *Mephitis mephitis* | 3 | Y | 25 | N | N | N |
| *Vulpes vulpes* |  |  | 17 | N | N | N |
| *Odocoileus virginianus* |  |  | 20 | N | N | N |
| *Lynx rufus* |  |  | 3 | N | N | N |
| *Sylvilagus floridanus* |  |  | 118 | Y | Y | N |
| *Marmota monax* |  |  | 31 | Y | Y | N |
| *Tamias striatus* |  |  | 12 | N | N | N |
| *Lasiurus borealis* |  |  | 12 | Y | N | Y |
| *Eptesicus fuscus* |  |  | 7 | N | N | N |
| *Ursus americanus* |  |  | 7 | N | N | N |
| *Sciurus niger* |  |  | 3 | N | N | N |
| *Blarina brevicada* |  |  | 3 | N | N | N |
| *Rattus rattus* |  |  | 2 | N | N | N |
| *Urocyon cinereoargenteus* |  |  | 2 | N | N | N |
| *Castor canadensis* |  |  | 2 | N | N | N |
| *Peromyscus sp.* |  |  | 2 | N | N | N |
| *Mustela vison* |  |  | 1 | N | N | N |
| *Microtus pennsylvanicus* |  |  | 1 | N | N | N |
| *Mus musculus* |  |  | 1 | N | N | N |

**Supplementary Table 9.** Comparison of results from this study, experimental infection studies and predicted susceptibility based on modeling of the ACE2 receptor. RT-qPCR and seroprevalence data from our study are combined with data from previously published research on whether species were capable of being infected in the lab and their predicted susceptibility based on modeling of the ACE2 receptor. For species that do not have an exact match we included closely related species, which are indicated after the semicolon. * Indicates instances where a species has not been evaluated but information for a closely related species is available.

| Common Name | RT-qPCR Prevalence^1^ | Seroprevalence^1^ | Experimental Infection | ACE2 modeling |
| --- | --- | --- | --- | --- |
| Deer mouse | 5% | 7% | Seroconverted, live virus isolation^3-5^ | High^5-8^, medium^9^ |
| Striped Skunk | 0% | 0% | Seroconverted^3,10^, live virus isolation^3,10^ | Very low (Western spotted skunk)^9^ |
| Raccoon | 5% | 36% | Seroconverted, no virus isolated^10^ | Low^6,8^ |
| White-footed mouse | 0% | 17% |  | High^5,7^, low^8^ |
| Grey squirrel | 0% | 57% | *No seroconversion or virus isolation (Fox squirrel)^3^ | *High (Red squirrel)^11^ |
| Virginia opossum | 3% | 38% |  | *High (Gray short-tailed opossum)^7^ |
| Eastern red bat | 8% | - | *Seroconverted, live virus isolation (Egyptian fruit bat^12^; Mexican free-tailed bat^13^. No seroconversion or virus isolation (Big brown bats)^14^ | *Low-high depending on bat species^7,9,15^ |
| Groundhog | 7% | - |  | *Low (Alpine marmot, Yellow-bellied marmot)^16^, medium (Alpine marmot)^7,9,16,17^ |
| White-tailed deer | 0% | - | Seroconverted, live virus isolation^18^ | High^6,9^, low^7^ |
| Eastern cottontail | 3% | - | *No seroconversion or virus isolation (European cottontail)^3^ | *Low (European rabbit)^6^, Medium (European rabbit)^9,16^ |
| Eastern chipmunk | 0% | - |  | *Low (Thirteen-lined ground squirrel)^9,16^, Medium (Daurian ground squirrel^9^, thirteen-lined ground squirrel^9^) |
| Red fox | 0% | - | Seroconverted, live virus isolation^19^ | Low^9^, medium^7^ |
| ^1^Prevalence calculated from data collected in this study. All samples were collected from wildlife in Virginia from May 2022-Sept 2023. RT-qPCR results were considered positive if they had at least two of the three genes tested as positive (N, E, and S with a Ct<40) | | | | |

**Supplementary Table 10.** Estimates of monthly human presence at the 5 sites we trapped and collected serological data.

| Site | County | Monthly Use Estimates | Human presence Group |
| --- | --- | --- | --- |
| Mountain Lake Biological Station | Giles | 350 | Low |
| Brush Mountain | Montgomery | 10 | Low |
| Blacksburg | Montgomery | 84,188 | High |
| Roanoke Parks | Roanoke | 79,324 | High |
| New River Trails State Park | Wythe | 6,949 | High |

**Supplementary Table 11.** Housekeeping primer sequences for RT-qPCR testing in 22 wildlife species for presence of SARS-CoV-2. See attached excel file.

| **Supplementary Table 12**. Average and maximum lifespan of wild animals based on published literature. | | |
| --- | --- | --- |
| Species | | Average lifespan in the wild (years); max is in parenthesis |
| *Procyon lotor* | 5^a^ (20^b^) | |
| *Didelphis virginiana* | 1.5-2^a^ | |
| *Mephitis mephitis* | <1^a^ (6^b^) | |
| *Sciurus carolinensis* | (12.5^b^) | |
| *Peromyscus maniculatus* | <1^a^ | |
| *Peromyscus leucopus* | 1^a^ | |
| *Blarina brevicauda* | (2.5^b^) | |
| *Marmota monax* | 4-6^a^ | |
| *Lasiurus borealis* |  | |
| *Sylvilagus floridanus* | <3^a^ (5^b^) | |
| *Tamias striatus* | <2^a^ (3^b^) | |
| *Vulpes vulpes* | 3^a^ (7^b^) | |

^a^ Myers, P., R. Espinosa, C. S. Parr, T. Jones, G. S. Hammond, and T. A. Dewey. 2024. The Animal Diversity Web (online). Accessed at <https://animaldiversity.org>

^b^Carey, J. R., & Judge, D. S. 2002. Longevity records: Life spans of mammals, birds, amphibians, reptiles, and fish. Monographs on population aging, (8).

**Supplementary Table 13**. Buffer width set for use to estimate imperviousness and population density. Buffer width differed by species to represent the space used by each individual trapped.

| Species | Buffer width (m) | Source |
| --- | --- | --- |
| *Peromyscus spp.* | 50 | Wolff, J. O. 1985. The effects of density, food, and interspecific interference on home range size in *Peromyscus leucopus* and *Peromyscus maniculatus*. Canadian Journal of Zoology 63:2657-2662. |
| *Sciurus carolinensis* | 225 | Koprowski, J. L., K. E. Munroe, and A. J. Edelman. 2016. Gray not grey: Ecology of Sciurus carolinensis in their native range in North America. The Grey Squirrel: ecology & management of an invasive species in Europe. Woodbridge, Suffolk UK: European Squirrel Initiative, 1-18. |
| *Didelphis virginiana* | 500 | Gallo, T., et al. 2022. Mammals adjust diel activity across gradients of urbanization. Elife 11:e74756. |
| *Mephitis mephitis* | 1000 | Gallo, T., et al. 2022. Mammals adjust diel activity across gradients of urbanization. Elife 11:e74756. |
| *Procyon lotor* | 1000 | Gallo, T., et al. 2022. Mammals adjust diel activity across gradients of urbanization. Elife 11:e74756. |

**Supplementary Table 14**. Summary of sequences used to assemble phylogenetic trees for comparing the SARS-CoV-2 sequences isolated from wild animals to those isolated from humans in our geographical region. All human sequences were sourced from NCBI and GISAID.

| Pango lineage | Geographical region | Collection date ranges | Non-duplicate sequences |
| --- | --- | --- | --- |
| BA.2.10.1 (21L) | KY, NC, TN, VA, WV | 01/01/2022 - 10/19/2023 | 58 |
| XBB.1.5.10 (23A) | KY, NC, TN, VA, WV | 06/01/2023 - 10/19/2023 | 59 |
| XBB.1.5.45 (23A) | North America | 01/01/2022 - 10/21/2023 | 14 |
| EG.5.1.1 (23F) | KY, NC, TN, VA, WV | 07/01/2023 - 09/01/2023 | 139 |
| JD.1 | North America | 01/01/2022 - 10/21/2023 | 11 |
| XBB (22F)* | KY, NC, TN, VA, WV | 09/01/2022 - 10/19/2023 | 53 |
| XBB.1.5 (23A) | KY, NC, TN, VA, WV | 07/01/2023 - 10/19/2023 | 109 |
| XBB.1.16 (23B) | KY, NC, TN, VA, WV | 06/01/2023 - 08/01/2023 | 99 |

*Only sequences from GISAID were included

**Supplementary Table 15.** Cross-reference tables for the eight phylogenetic trees (Fig 3 & Supplementary Figures 2-9). See attached.
